# Kinetics of local C3 production orchestrates neutrophil recruitment in lung injury

**DOI:** 10.64898/2026.02.17.706326

**Authors:** Rafael Aponte Alburquerque, Josue I. Hernandez, Aasritha Nallapu, Marick Starick, Neelou Etesami, Sandip K. Mukherjee, Ayse Naz Ozanturk, Jose Vazquez, Allison Chang, Ashley Zheng, Lorena Garnica, Alberto Lopez, Andrew E. Gelman, John Belperio, Jae Woo Lee, Devesha H. Kulkarni, Alexander Hoffmann, Hrishikesh S. Kulkarni

**Author notes:** **Corresponding Author:** Hrishikesh Kulkarni, MD, MSCI University of California Los Angeles, 650 Charles E Young Drive S, Los Angeles, CA 90095.

## Abstract

Complement component 3 (C3) is crucial for host defense against bacteria. While the liver is the primary source of circulating C3, local C3 production at barrier surfaces such as the lung is key in early responses. Yet, how local complement-mediated responses are initiated at mucosal barriers is unknown. This study investigates the kinetics and necessity of lung-derived C3 during the initial hours of an infection. Using models of bacterial pneumonia in ex vivo-perfused human lungs and mice deficient in liver-derived C3, we demonstrate that intrapulmonary C3 production and activation precedes the accumulation of circulating C3 into the bronchoalveolar space. Utilizing mice deficient in lung-derived C3, we demonstrate that epithelial cell-derived C3 is required for early neutrophil recruitment in pneumonia. Transcriptomic and proteomic analyses reveal that neutrophil chemotactic pathways such as C5a and CXCL2 depend on lung epithelial cell-derived C3. These findings demonstrate how lung epithelial-derived C3 influences early mucosal responses to infection via both canonical (direct) and non-canonical (indirect) pathways.

**SUMMARY:** Alburquerque et al show an initial, entirely local phase of complement-mediated mucosal protection before a subsequent, systemic response occurs in the setting of barrier disruption. Their work suggests that complement component C3 derived locally at a barrier from the epithelium influences early responses to infection by recruiting neutrophils via multiple pathways independent of circulating C3.

## INTRODUCTION

Barrier surfaces lined by epithelial cells form the first line of host defense in infection (Nowarski et al., 2017). Bacterial infections of the lungs can result in pneumonia, and when uncontrolled, in acute lung injury (ALI), which is characterized by exuberant inflammation, alveolar-capillary barrier disruption, and impairment of gas exchange (Kulkarni et al., 2022; Traber and Mizgerd, 2025). Epithelial cells produce host defense proteins such as cytokines, chemokines, and antimicrobial peptides early during infection, to both limit pathogen burden and recruit immune cells to the site of injury (Leiva-Juárez et al., 2018; Hewitt and Lloyd, 2021). These initial responses are followed by the entry of circulating host defense proteins into the site of infection (Medzhitov, 2007). Among these proteins, components of the complement system are becoming increasingly recognized as key players in the local response to infection and injury at mucosal barriers such as the skin, gut, and lung (Kolev et al., 2020; Sahu et al., 2023; Wu et al., 2024).

C3 is a central component of the complement system and plays a critical role in immune defense against pathogens, including those affecting the respiratory tract (Quezado et al., 1994; Mueller-Ortiz et al., 2004; Rattan et al., 2017). C3 cleavage facilitates amplification of the complement cascade, leading to C5 cleavage, the generation of anaphylatoxins and opsonins, and the formation of membrane attack complexes on pathogen surfaces (Sahu et al., 2022; Kulkarni et al., 2024). These anaphylatoxins facilitate rapid chemotaxis of phagocytes into the lungs, and clearance of pathogens (Kazmierowski et al., 1977; Neupane et al., 2020). We and others have shown that C3 increases at the pulmonary mucosal barrier in response to infection and injury (Bolger et al., 2007; Sahu et al., 2023). The majority of circulating C3, found at concentrations of 1-2 mg/ml in humans, is produced by the liver (Torisu et al., 1972). This circulating, liver-derived C3 leaks into the bronchoalveolar space during ALI (Bolger et al., 2007). However, we previously demonstrated that this accumulation takes time, and lung-derived C3 production takes precedence over liver-derived C3 in protecting the host against severe bacterial pneumonia (Sahu et al., 2023).

Cellular sourcing of C3 in the lungs primarily occurs from cells of the mesothelial/fibroblast, macrophage and epithelial lineage (Chaudhary et al., 2022). We and others have previously reported that C3 can be synthesized and secreted into the airspace by lung epithelial cells (Kulkarni et al., 2019; Yan et al., 2021). Moreover, C3 expression increases over three-fold in epithelial cells forming the barrier surface at 24 h post-bacterial pneumonia (Sahu et al., 2023). This local increase in C3 levels and its activity in the bronchoalveolar lavage fluid (BAL) occurs within the first 24 h of infection with *Pseudomonas aeruginosa* (*Pa*) and mitigates the severity of pneumonia-induced ALI independent of circulating C3 levels (Sahu et al., 2023). The absence of lung epithelial cell-derived C3 resulted in increased epithelial cell injury and loss, as well as increased lung inflammation and alveolar-capillary barrier disruption, independent of bacterial proliferation (Sahu et al., 2023). A tissue-specific complement response also occurs in the setting of viral pneumonia, and precedes lung injury and inflammation (Szachowicz et al., 2025). However, the precise kinetics and mechanisms through which lung epithelial cell-derived C3 initiates an early local immune response following infection, remain unclear.

Given the delay in liver-derived C3 entering the lungs, this study aimed to investigate the initiation of lung-derived C3 production and activation early during a bronchopulmonary infection and its role in triggering a local host immune response.

## RESULTS

### Accumulation and cleavage of C3 increases at 4 h of bronchopulmonary infection

To investigate how complement is triggered at a mucosal barrier surface, we leveraged an *ex vivo* human lung model infected with *Pa* and a mouse model of *Pa* pneumonia-induced ALI. In the mouse model (**Figure 1A**), BAL levels of C3 and its activation products, as measured by their ability to deposit on lipopolysaccharide (LPS), rose after 4 h of infection (**Figure 1B**). On the contrary, circulating C3 levels remained stable over time (**Figure 1C**). The results from our LPS assay were also consistent with a customized ELISA for detecting intact/full-length C3 levels (**Supplementary Figures S1A & B**), which demonstrated, on average, 263 ng/mL of C3 in the BAL of uninfected mice that tripled by 4 h and increased six-fold by 8 h post-*Pa* infection (**Figure 1D**). To measure local activation of C3 in the mouse lung, we adapted a previously reported protocol for measuring circulating C3a levels (Pagano et al., 2009) to detect a neo-epitope on C3a in the BAL (**Supplementary Figures S1C & D**). This neo-epitope is only detected when C3a is cleaved from C3 and is thus, a direct measure of C3 activation. C3a levels began to increase at 2 h post-infection, reaching a significant peak at 4 h and doubling at 8 h (**Figure 1E**). To investigate this hypothesis in humans, we used an ex vivo model of perfused human lung (**Figure 1F**) (Lee et al., 2009). Both C3 and its activation fragments (C3b/iC3b) also increased in the BAL at 4 h post-*Pa* in the ex vivo human lung model compared to non-infected lungs. C3 activation was also detected in the absence of any blood in the perfusate (**Figures 1G & H**). To assess if this local C3 activation was generated by circulating C3 entering the airspaces, we analyzed BAL IgM levels, a measure of alveolar-capillary barrier disruption (Kulkarni et al., 2022), which did not change between 0 and 4 h post-*Pa* (**Figure 1I**). This localized accumulation and activation in both mice and humans suggests that the lung is generating its own immune response during the initial stages of infection. These observations are novel in demonstrating that local C3 sourcing is key for pathogen recognition and immune activation in response to a bacterial infection.

**Figure 1.**
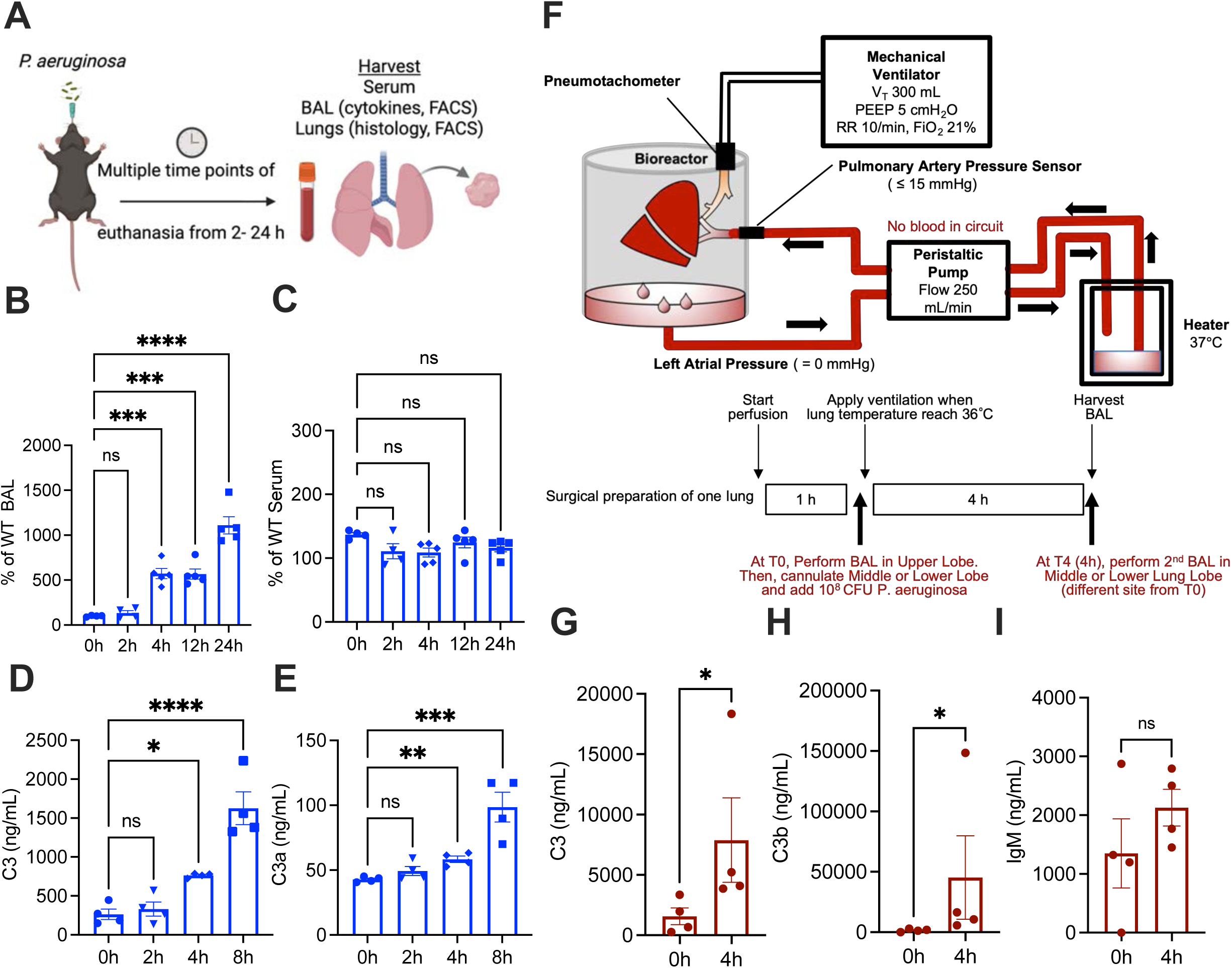
Production and cleavage of C3 starts to increase at 4 h post bronchopulmonary *Pa* infection. (**A**) Schematic of infection in mouse model. Created in BioRender. Kulkarni, H. (2026) https://BioRender.com/jevlom7 (**B-C**) Assay for detecting C3 activation on lipopolysaccharide (LPS) normalized to wildtype (WT) uninfected (**B**) bronchoalveolar lavage (BAL) and (**C**) serum of WT mice infected with *Pseudomonas* (Pa) and euthanized at 2, 4, 8, 12 and 24 h. Results for (**D**) full-length C3 ELISA and (**E**) C3a-neoepitope ELISA in BAL post-*Pa* infection. (**F**) Schematic of infection in ex vivo human lung model. Modified based on Zhang et al. (**G-I**) Changes in BAL (**G**) C3, (**H**) C3b/iC3b, and (**I**) IgM levels in ex vivo human lung pre-infection (T0, 0 h) and post-infection (T4, 4h). Distribution of data is represented as scatterplots showing individual data points, and bars represent means ± SD. ****P<0.0001, ***P < 0.001, **P < 0.01, and *P < 0.05 using ordinary one-way ANOVA after adjusting for multiple group comparisons, and Mann-Whitney test for two-group comparison. Abbreviations: CFU: colony-forming units; FACS: fluorescence-associated cell sorting (flow cytometry); PEEP: positive end-expiratory pressure; RR: respiratory rate; VT: tidal volume.

### Production and activation of C3 in the lungs precedes leakage of circulating C3 into the bronchoalveolar space

Since the liver is the major source of circulating C3 in an intact *in vivo* system (Torisu et al., 1972), we wanted to eliminate this source for investigating the kinetics of lung-derived C3. Hence, we used a liver-derived C3-deficient mouse model (*C3^f/f^ AlbCre^+^*) (Sahu et al., 2023). Despite the absence of any circulating C3 (**Supplementary Figure S1E**), full-length C3 increased in the BAL of these liver-derived-C3-deficient mice starting at 4 h after *Pa* infection (**Figure 2A**). Additionally, C3a levels (**Figure 2B**) and its deposition on LPS (**Figure 2C**) were also significantly increased at 4 h after infection. The early rise in BAL C3 and C3a levels, even in the absence of circulating C3, underscores the lung’s ability to induce a localized complement response, independent of systemic contributions within the initial hours of infection.

**Figure 2.**
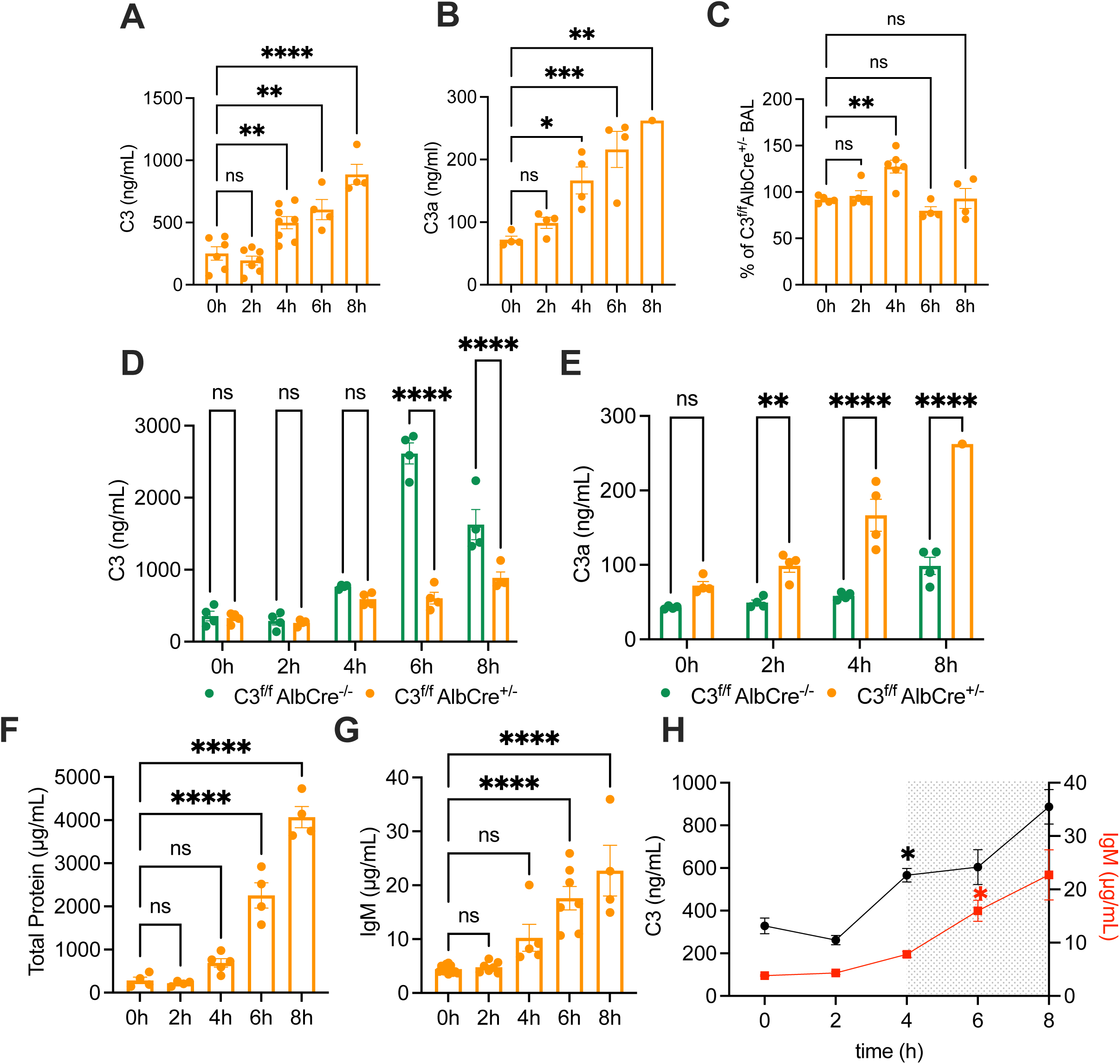
Local production and activation of C3 starts before entry of circulating C3 into the lungs. Full-length ELISA measuring (**A**) C3 and (**B**) C3a in BAL of C3^f/f^ AlbCre mice euthanized from 0-8 hours after *Pa* infection. (**C**) LPS assay on BAL of infected C3 liver-deficient mice, normalized to activity of uninfected BAL. Full-length ELISA for (**D**) C3 and (**E**) C3a in BAL comparing C3^f/f^AlbCre mice (orange) and littermate controls (green). (**F**) Total Protein by BCA assay and (**G**) IgM were performed on BAL. (**H**) Representative comparison between levels of IgM and C3 in BAL of C3^f/f^ AlbCre mice over time. Distribution of data is represented as scatterplots showing individual data points, and bars represent means ± SD. ****P<0.0001, ***P < 0.001, **P < 0.01, and *P < 0.05 using unpaired two-tailed t test.

To determine whether the liver contributes to lung C3 levels, we compared *C3^f/f^AlbCre^+^* with littermate *C3^f/f^*controls. We found that the initial increase in BAL C3 and C3a levels similarly occurred in in C3^f/f^ controls that had intact circulating C3 levels. However, at 6 h post-infection, the control mice showed a significant increase in BAL C3 levels over the *C3^f/f^AlbCre^+^*mice, suggesting that this was the time point at which circulating C3 starts accumulating in the lungs **(Figure 2D**). Moreover, BAL C3a were significantly higher at 2 h post infection in *C3^f/f^AlbCre^+^*mice, suggesting that local C3 activation occurs prior to circulating C3 entering the lungs (**Figure 2E**). Aligning with these observations, BAL total protein levels were also significantly higher at 6 h post-infection in both liver-derived C3 deficient mice (**Figure 2F**) and their littermate controls (**Supplementary Figure S1F**), suggesting a disruption of the alveolar-capillary barrier. To confirm this observation, we measured IgM levels in the BAL. In line with total protein levels, BAL IgM increased significantly starting at 6 h post-infection compared to baseline IgM levels (**Figure 2G**). When changes in the BAL C3 levels were compared to changes in BAL IgM levels in the liver-derived-C3-deficient mice, the increase in BAL C3 levels occurred prior to the increase in BAL IgM levels (**Figure 2H**). These results, along with the results from our ex vivo lung model, suggest that a local increase in C3 production in the lung occurs prior to leakage of circulating C3 into the bronchoalveolar space (**Figure 2H**). These findings suggest that C3 is produced early in response to an infection before circulating C3 can reach the lungs to facilitate pulmonary host defense.

BAL total protein levels were also significantly higher at 6 h post-infection in both C3 liver-deficient mice (**Figure 2F**) and their littermate controls (**Supplementary Figure S1F**), suggesting that a disruption of the alveolar-capillary barrier is observed at this timepoint in our model. To confirm this observation, we measured IgM levels in the BAL. Similar to the total protein levels, BAL IgM was significantly higher starting at 6 h post-infection compared to baseline IgM levels (**Figure 2G**). When changes in the BAL C3 levels were compared to those in BAL IgM levels in the liver-deficient C3 mice, the increase in BAL C3 levels occurred prior to the increase in BAL IgM levels (**Figure 2H**). These results, along with the results from our *ex vivo* human lung model, suggest that a local increase in C3 production in the lung occurs before the leakage of circulating C3 into the bronchoalveolar space (**Figure 2H**). These observations lead us to speculate that complement-mediated mucosal host defense against an acute insult comprises of an entirely local response comprising of early C3 production and activation. This early local phase is followed by a systemic phase when circulating components later reaching the site after alveolar-capillary barrier disruption to sustain host defense and restore mucosal homeostasis.

### Initial pulmonary neutrophil recruitment during bronchopulmonary *Pa* infection occurs independent of liver-derived C3

To examine the physiological importance of early local C3 activation, we evaluated the cellular immune response within the early timepoints of a bronchopulmonary *Pa* infection. We performed flow cytometry on the lung digest and BAL pellet from liver-derived-C3-deficient mice. Intrapulmonary CD45^+^ immune cells started to increase as early as 2 h post-infection. Additionally, leukocyte accumulation was significantly higher at 4 h after infection compared to uninfected mice, even in the absence of any circulating C3 (**Figure 3A**). Analysis of intrapulmonary leukocyte phenotypes at 4 h post-infection revealed that most cells were CD45^+^Ly6g^+^ neutrophils (**Figure 3B**).

**Figure 3.**
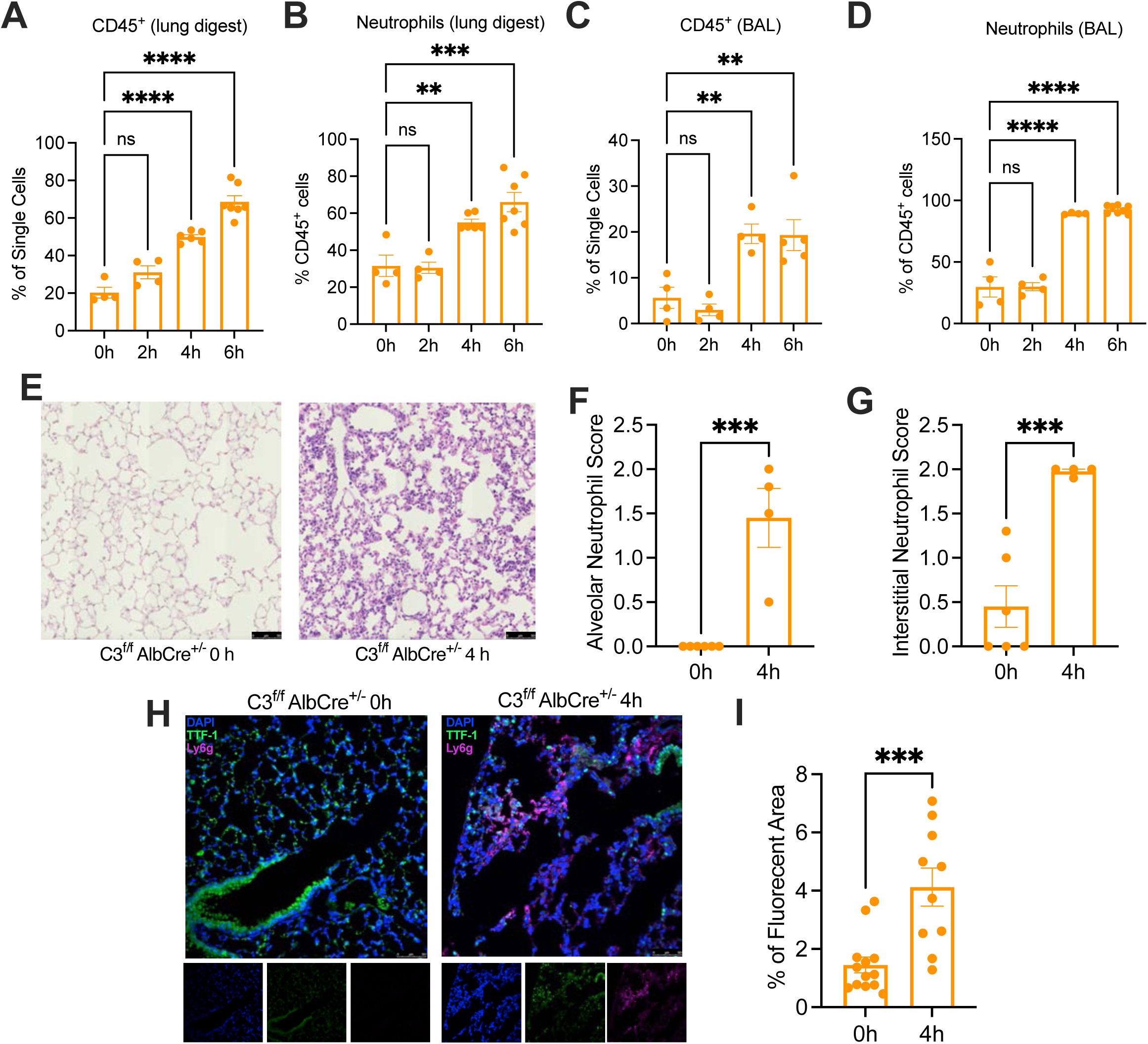
Pulmonary neutrophil recruitment occurs within the initial hours of a bronchopulmonary infection independent of circulating C3 levels. Cell population frequencies for (**A**) CD45^+^ cells and (**B**) neutrophils (CD45^+^, Ly6g^+^) in whole lung and in (**C & D**) BAL were measured by flow cytometry. (**E**) Representative lung histology using H&E staining of alveolar space (n=4). Lung scoring for (**F)** alveolar and (**G**) interstitial neutrophils. (**H**) Immunofluorescence staining for neutrophils (Ly6g) and lung epithelial cells (TTF-1). (**I**) Percentage of fluorescent area for neutrophils. Distribution of data is represented as scatterplots showing individual data points, and bars represent means ± SD. ****P < 0.0001, ***P < 0.001, **P < 0.01, and *P < 0.05, unpaired two-tailed t test.

Neutrophils are highly responsive immune cells that mobilize to sites of infection or tissue injury (Mócsai, 2013; Peiseler and Kubes, 2019). Upon detecting bacterial infection, they rapidly transmigrate across the vascular endothelium from the bloodstream into affected tissues typically within a few hours (Nourshargh and Alon, 2014). In our *Pa* infection model, we observed an increase in BAL CD45^+^ cells, primarily neutrophils, within 4 h of infection in liver-derived-C3-deficient mice, demonstrating that these immune cells had entered the lumen independent of any circulating C3 (**Figures 3C & D**). Histological and immunofluorescent analysis demonstrated increased neutrophil infiltration in both the alveolar and interstitial spaces compared to uninfected lungs (**Figures 3E-I, Supplementary Figures S2A-D**).

### Early pulmonary neutrophil recruitment during pneumonia is dependent on lung epithelial cell-derived C3

We noted that intrapulmonary neutrophil recruitment preceded the increase in total protein and IgM in the BAL, suggesting a response to local C3 production and activation. Hence, to determine whether neutrophil recruitment depends on the initial lung-derived C3 activity, we sought a mouse model that is defective in local lung C3 production. The *C3^f/f^Scgb1a1-CreER^T2^* mouse allows for tamoxifen-dependent knockout of C3 in epithelial cells, which can then be monitored by tdTomato expression (Sahu et al., 2023). We infected *C3^f/f^Scgb1a1-CreER^T2^*mice following tamoxifen treatment or control with *Pa*. We found a significantly lower proportion of tdTomato^+^ epithelial cells in tamoxifen-treated club-cell-C3-deficient mice compared to similarly treated controls at 4 h post*-Pa* infection (**Figure 4A**). Moreover, BAL levels of both intact C3 (**Figure 4B**) as well as cleaved C3a (**Figure 4C**) were lower in infected club-cell-C3-deficient mice compared to their controls at 4 h post-*Pa* infection. However, the BAL levels of chemokines CXCL1 (**Figure 4D**) and CXCL2 (**Figure 4E**) were also lower in the club-cell-C3-deficient mice at 4 h post-*Pa* infection compared to their controls. In contrast, the levels of other cytokines, such as TNF-α and IL-1β were not significantly different (**Figures 4F & G, Supplementary Figure S2E-H**). In parallel, lung neutrophilia was significantly lower in club-cell--C3-deficient mice at 4 h post-infection as compared to their controls (**Figure 4H**). Immunofluorescent analysis also revealed fewer intrapulmonary neutrophils in club-cell-C3-deficient mice as compared to their littermate controls (**Figures 4I&J**) These data suggest lung epithelial cell-derived C3 facilitates neutrophil recruitment in the initial hours of a bacterial pneumonia.

**Figure 4:**
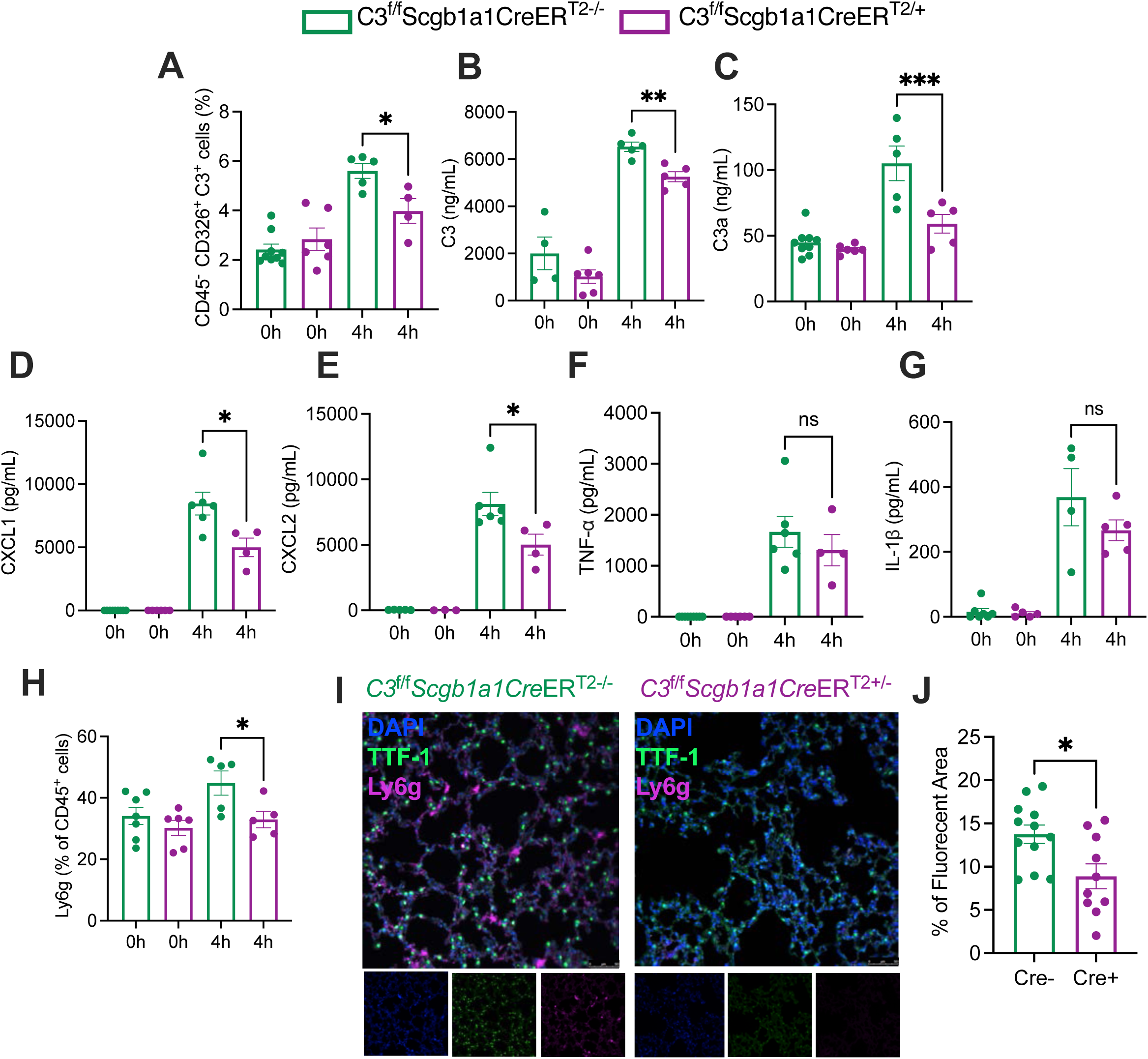
C3-deficient epithelial cells result in diminished neutrophil recruitment at initial hours of *a* bronchopulmonary infection. Cell population frequencies for (**A**) percentage of C3^+^ epithelial cells via flow cytometry (CD45^+^, CD326^+^) in the lungs of C3^f/f^Scgb1a1CreER^T2+/-^ (purple) and C3^f/f^Scgb1a1CreER^T2-/-^ (green) mice at 4 h post infection. (**B**) ELISAs for Full-length C3 ELISA and (**C**) for C3a in BAL. Multiplex ELISA for (**D**) CXCL1, (**E**) CXCL2, (**F**) TNF-α and (**G**) IL-1β. (**H**) Percentage of Neutrophils (CD45^+^, Ly6g^+^) in whole lung measured by flow cytometry. (**I**) Immunofluorescence staining for neutrophils (Ly6g) and lung epithelial cells (TTF-1). (**J**) Percentage of fluorescent area for neutrophils. Distribution of data is represented as scatterplots showing individual data points, and bars represent means ± SD. ***P < 0.001, **P < 0.01, and *P < 0.05 using unpaired two-tailed t test.

### Lung epithelial cell-derived C3 promotes intrapulmonary neutrophilia via both canonical and non-canonical mechanisms

To investigate how lung epithelial cell-derived C3 facilitates pulmonary neutrophil recruitment, we first performed bulk RNA sequencing on lungs from *Pa*-infected mice deficient in lung epithelial cell-derived C3 at 4 h post-infection, as well as corresponding controls (**Figure 5A**). Over 100 genes were downregulated in the lungs of the *C3^f/f^Scgb1a1-CreER^T2^*mice as compared to their controls, including genes such as *Cxcl1*, *Cxcl2* and *C5ar1* among others (**Figure 5A, Supplementary Figure S3A**). Enrichment analysis of the downregulated genes identified “Inflammatory Response”, “Granulocyte Chemotaxis”, and “Neutrophil Migration” as the top three ontologies (**Figure 5B**). COMPBIO—a biological knowledge generation platform that employs contextual language processing algorithms to identify enriched core biological concepts associated with these downregulated genes utilizing an ontology-free approach (Liu et al., 2021; Silver et al., 2023)—identified the top three themes as “Myeloid Infection Response/Interferon Signaling,” “N-formyl Peptide Receptor,” and “C5a Anaphylatoxin Chemotactic Receptor” (**Figure 5C**). 18 genes were identified common to these three themes, which included *Cxcl1*, *Cxcl2 and C5ar1* (**Figure 5C**).

**Figure 5.**
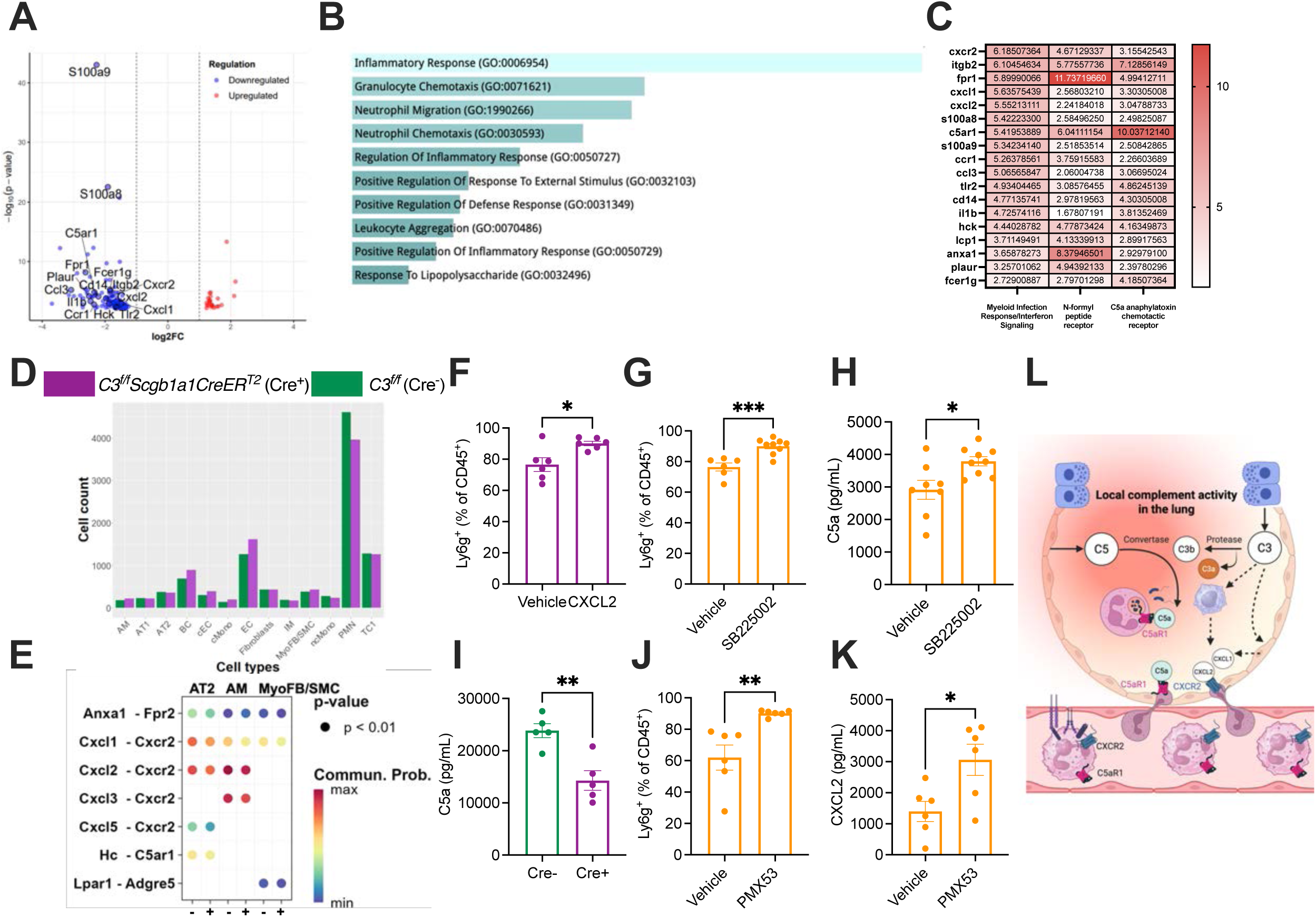
Neutrophil recruitment to the lung in the initial hours of a bronchopulmonary infection occurs through C3-dependent direct and indirect pathways. (A-C) Bulk RNA sequencing (RNA-seq) was performed on whole lung. (**A**) Volcano plot comparing genes obtained from differential analysis expression and filtered by FDR step up ≤0.05 in lungs of mice deficient in epithelial cell-derived C3 versus controls at 4 h post-*Pa* infection. (**B**) EnrichR analysis of the 130 genes from Figure S3A. (**C**) Table with the genes involved in the top 3 significant themes and organized by their enrichment score determined by COMPBIO. (**D**) Proportion of cells on single cell RNA-Seq of lungs from *C3^f/f^Scgb1a1CreER^T2^* mice (Cre^+^, purple) and littermate controls (Cre^-^, green) at 4 h post infection (n=3 mice/group). (**E**) CellChat analysis of (D). AT2: Type 2 alveolar epithelial cells; AM: alveolar macrophages; MyoFB/SMC: myofibroblasts/smooth muscle cells. (**F**) Neutrophils in BAL of *C3^f/f^Scgb1a1CreER^T2^* mice given CXCL2 orotracheally, infected with *Pa* and euthanized at 4 h post-infection. (**G & H**) Liver-deficient C3 mice (C3^f/f^AlbCre, orange) injected intraperitoneally (IP) with SB225002, a CXCR2 antagonist, infected with *Pa* and euthanized at 4 h post-infection. Neutrophils (**G**) and C5a levels (**H**) measured in BAL. (**I**) BAL C5a levels in *C3^f/f^Scgb1a1CreER^T2^* mice (purple) and controls (green) mice at 4 h post-*Pa* infection. (**J & K**) Liver-deficient C3 mice (C3^f/f^AlbCre, orange) orotracheally administered PMX53, a C5aR1 antagonist, infected with *Pa* and euthanized at 4 h post-infection. Neutrophils (**J**) and CXCL2 levels (**K**) measured in BAL. Data distribution is represented as scatterplots showing individual data points, and bars represent means ± SD. *P < 0.05, **P < 0.01, ***P<0.001 using unpaired two-tailed t test. (**L**) Schematic proposal of how lung epithelial cell-derived C3 promotes intrapulmonary neutrophilia in the initial hours of an infection. Created in BioRender. Kulkarni, H. (2026) https://BioRender.com/2l07l29

To investigate the cellular communication networks altered by lung epithelial cell-derived C3 in the setting of an acute bacterial pneumonia, we performed single cell RNA-sequencing (scRNA-seq) on the lungs of *C3^f/f^Scgb1a1-CreER^T2^*mice at 4 h post-*Pa* infection and compared the signatures to similarly treated littermate controls. Aligning with our flow cytometry and immunofluorescence data, mice deficient in lung epithelial cell-derived C3 exhibited lesser intrapulmonary neutrophilia at 4 h post-*Pa* infection compared to controls (**Figure 5D**). Moreover, CellChat-derived intercellular communication networks identified several chemotactic pathways downregulated in club cell-deficient C3 mice compared to controls (**Figure 5E**). Given that gene and protein levels of CXCL2 were lower in the lungs of *Pa*-infected mice deficient in lung epithelial cell-derived C3 at 4 h post-infection (**Figures 4E & 5A**) and CXCL2 is a more potent chemokine than CXCL1 in certain contexts (Sawant et al., 2021), we subsequently assessed if epithelial cell-derived C3 recruited pulmonary neutrophils via CXCL2. Intratracheal instillation of CXCL2 in mice deficient in epithelial cell-derived C3 overcame the defect in neutrophil recruitment into the bronchoalveolar space at 4 h post-*Pa* infection (**Figures 5F**). The lungs of these mice demonstrated an upregulation of transcripts such as *Fpr2*, *Il17a* and, *Il22* at 4 h post-*Pa* infection (**Supplementary Figures 3B & C**), which have been implicated in neutrophil recruitment to sites of inflammation (Ye et al., 2001; Yaeger et al., 2025).

To determine if lung-derived C3 facilitates neutrophil recruitment via the CXCL2-CXCR2 axis, we administered a CXCR2 antagonist (SB225002) to liver-derived-C3 deficient mice (*C3^f/f^AlbCre)*, who have intact intrapulmonary C3 production and activation (**Figure 2**). Although we observed a decrease in BAL neutrophilia in wildtype mice in the setting of CXCR2 antagonism (**Supplementary Figure S3D**), blocking CXCR2 in *C3^f/f^AlbCre* mice resulted in a paradoxical increase in BAL neutrophilia (**Figure 5G & Supplementary Figure S3E**). Coincidentally, BAL C5a levels were increased in these mice the setting of CXCR2 antagonism (**Figure 5H**). Aligning with these observations, BAL C5a levels were lower in *Pa*-infected mice deficient in lung epithelial cell-derived C3 at 4 h post-infection compared to their controls (**Figure 5I**), suggesting that lung epithelial cell-derived C3 influences local C5a levels. However, orotracheal administration of PMX53, a C5aR1 antagonist, resulting in a paradoxical increase in BAL neutrophilia (**Figure 5J & Supplementary Figures S3G-I**) and CXCL2 (**Figure 5K**) independent of bacterial burden (**Supplementary Figure S3J**). These observations suggest that lung epithelial cell-derived C3 promotes neutrophil recruitment not only via C5a, a direct, canonical effector of complement cascade activation, but also indirectly via chemokines such as CXCL2.

### Orchestration of an early immune response by C3 derived from the lung barrier surface

Taken together, our findings indicate that C3 produced and activated in the lungs is key for initiating an immune response during the early hours of an infection even before C3 from the circulation can accumulate in the lung. Thus, these data suggest an initial, entirely local phase of complement-mediated mucosal protection before a subsequent, systemic response occurs in the setting of alveolar-capillary barrier disruption. We have previously reported the necessity of lung epithelial-derived C3 in host defense against bacterial pneumonia-induced lung injury (Sahu et al., 2023). Yet, how this epithelial cell-derived C3 is operative at barrier surfaces has been unclear. Our new findings reveal that lung epithelial cell-derived C3 is required for neutrophil recruitment into the lungs during the early hours of an insult to promote mucosal host defense. The mechanisms for this recruitment are multifactorial, and involve both direct effects of complement activation (e.g. C5a generation) and indirect effects (e.g., via CXCL2) (**Figure 5L**).

C5 is locally produced in the lung by epithelial cells (Posch et al., 2021; Chaudhary et al., 2022) and is cleaved to C5a upon activation, which binds to C5aR1 on neutrophils to recruit them (Bosmann and Ward, 2012; Kumar et al., 2020). CXCL1 and CXCL2 are produced by epithelial cells, macrophages, and endothelial cells, and are known chemoattractants for neutrophils (Prame Kumar et al., 2018; Lentini et al., 2020). On one hand, our data would suggest that C3 production and activation facilitates the amplification of the complement cascade and the cleavage of C5 to C5a, a known chemoattractant (Bosmann and Ward, 2012). On the other hand, we also show that C3 from lung epithelial cells influences the production of chemokines such as CXCL2, which are key to neutrophil recruitment. Thus, our data indicate that lung epithelial cell-derived C3 promotes neutrophil recruitment via multiple ligand-receptor interactions rather than a single regulatory axis (**Figure 5L**). These observations are supported by reports from other models such as lung cancer, which suggest that neutrophil recruitment into the lungs may not occur through a single axis (Kwak et al., 2024). The cell types and mechanism by which lung epithelial cell-derived C3 induces these chemokines is currently under investigation (Earhart et al., 2024). Interestingly, prior work in a murine model of dermal *Leishmania* infection suggest that neutrophils do not respond to IL-1β or macrophage-derived molecules, but are rather guided by the C5a-C5aR1 axis, using L-selectin and integrins, to extravasate into the draining lymph node parenchyma (Prat-Luri et al., 2022). These findings unravel the novel and crucial role of barrier-sourced complement in recruiting neutrophils early in an infection.

A limitation of our study is that although the *C3^f/f^Scgb1a1-CreER^T2^* mice are useful to assess the dependence of immune responses on lung epithelial cell-derived C3, an epithelial cell-specific C3-overexpressing mouse would be more helpful in assessing the effects of receptor antagonism. Additionally, our focus was on how lung epithelial cell-derived C3 influences neutrophil recruitment. As a result, we did not assess neutrophil functions such as oxidative burst, phagocytosis or NETosis, which play important roles in host defense after neutrophil recruitment (Lehman and Segal, 2020; Burgener and Schroder, 2020). We also did not test if the effects of C5a depend on endothelial C5aR2, which has been shown to transport C5a into the vessel lumen to initiate C5aR1-mediated neutrophil arrest in a murine model of immune complex-induced arthritis (Miyabe et al., 2019). However, utilizing a system where there is no circulating C3 but intact pulmonary C3, combined with interrogating the bronchoalveolar space at multiple time points using orthogonal approaches in both human and mouse lungs, we systematically assess complement-mediated defense at a surface such as the lung that constantly interacts with the environment. Our findings reveal a critical role for epithelial cell-derived C3 in the initiation of mucosal immune responses to an acute infection and set a precedent for investigating its impact at multiple barrier surfaces such as the skin, gut, and genitourinary tract.

## METHODS

### Experimental Design

The ARRIVE (Animal Research: Reporting of In Vivo Experiments) guidelines were followed for reporting of in vivo experiments, and are reported throughout the Methods and Figure Legends. Female and male mice between the age of 8 and 12 weeks of age were used in these experiments and were age-matched in each experiment. Sample size was determined based on prior studies on acute lung injury (Sahu et al., 2023). Pre-established exclusion criteria for mice included being pregnant or injured (i.e., unanticipated pre-existing wounds from co-housing). Each experiment was repeated at least twice; both biological and technical replicates were included. Observed effects were attributed to differences between groups only when specified.

### Mice

All animal experiments were conducted in accordance with protocols approved by the Institutional Animal Care and Use Committee at Washington University School of Medicine (WUSM) and the Division of Laboratory Animal Medicine (DLAM) at University of California Los Angeles (UCLA). Mice, aged 8 to 12 weeks and on a C57BL/6 background, were used throughout. C57BL/6J (RRID:IMSR_JAX:000664, termed wild-type, or WT) and B6.129S4-C3tm1Crr/J (RRID:IMSR_JAX:003641, termed C3-deficient, or C3KO) mice were obtained from Jackson Laboratory (Bar Harbor, ME, USA). The mice were maintained as groups in our animal housing facility at the Washington University in St. Louis School of Medicine with a 12 h light-dark cycle, with temperatures between 22-23°C and humidity of 68-72%. Genetically modified strains on a B6 background previously reported by our lab (including *C3^f/f^AlbCre* and *C3^f/f^ Scgb1a1-CreER^T2^* mice, and littermate controls, (Sahu et al., 2023)) were also maintained in our colonies. Genotyping was performed at Transnetyx (Cordova, TN), using a qPCR based system to detect the presence or absence of a target sequence within each sample.

Details on control groups, sample sizes, and experimental units are provided in each figure legend. While specific randomization was not used, efforts to minimize confounding included alternating infection and euthanasia between groups (e.g., Group 1, mouse 1; Group 2, mouse 1; Group 1, mouse 2; Group 2, mouse 2, etc.). All personnel involved in data collection were blinded to group allocations. All figures have been represented as scatter plots with individual data points, and range is represented as mean ± standard deviation.

### Preparation of bacteria for infection

To induce an immune response in our murine model, *Pseudomonas aeruginosa* (Pa) strain 57-15, a clinically relevant isolate from the sputum of a cystic fibrosis patient, was cultured and prepared as previously described (Sahu et al., 2023). Bacteria were cultured on LB agar plates (Sigma-Aldrich, #L7025-500TAB) for 16 h at 37°C from a frozen stock. The following day, *Pa* colonies were inoculated into LB broth (Sigma-Aldrich, #L3522) and incubated overnight at 37°C in a rotating shaker set to 220 rpm. After incubation, the bacteria were harvested by centrifugation at 3000g for 6 min at 4°C, and the resulting pellet was resuspended in sterile phosphate-buffered saline (PBS). The bacterial concentration was estimated by measuring absorbance at 380 nm using an Epoch Microplate Spectrophotometer (BioTek) to achieve the target dose of 2 × 10^5^ CFU/g of body weight.

### *In vivo* bacterial infection

Mice were infected with *Pseudomonas aeruginosa* (Pa) for 2, 4, 6, 8, 12, and 24 hours at a dose of 2 × 10^5^ CFU/g of body weight. Prior to infection, mice were anesthetized with an intraperitoneal injection of a ketamine (100 mg/kg) and xylazine (10 mg/kg) cocktail, followed by orotracheal intubaton. A fiber-optic cable connected to a light source (BioLITE Intubation System, BIO-MI-KIT, Braintree Scientific) illuminated the trachea, allowing precise placement of a 20-gauge Surflo catheter (#SR-OX2025CA, Terumo Medical Products). A suspension of *Pa* in sterile PBS was then administered through the catheter at a volume of 1 μl/g of body weight.

Euthanasia was conducted using Avertin, prepared as a 2% working solution from a stock solution (100% Avertin made by dissolving 1 g of 2,2,2-tribromoethyl alcohol (#T48402, Sigma-Aldrich) in 1 ml of tert-amyl alcohol (#240486, Sigma-Aldrich)). Following euthanasia, lungs were harvested for bronchoalveolar lavage (BAL) collection, flow cytometry analysis, or whole-organ fixation in 10% formalin for histological examination.

For BAL collection, 0.5 ml of PBS containing 1X Halt Protease Inhibitor Cocktail (diluted from 100X; #78429, Thermo Fisher Scientific) was instilled per lung and then recovered. The BAL fluid was centrifuged at 500g for 5 min to pellet cells, and the supernatant was collected and stored at −80°C for subsequent analyses.

### *Ex vivo* human lung model

For the ex vivo perfused human lung injured with *Pseudomonas aeruginosa* pneumonia (Zhang et al., 2021), deceased donor lungs (IRB-23-1362-CR-002, donated from One Legacy Donor Network, Azusa, CA) were perfused with Normal Saline (NS) containing 5% bovine serum albumin (BSA) (volume = 1350 ml) with a cardiac output of 0.25 L/min and then ventilated (Harvard Apparatus, tidal volume 300 ml, respiratory rate = 10/min, FiO2 = 21%, and PEEP = 5 cmH_2_O). Either the right or left lung was used. The upper lobe was first cannulated and a BAL was performed using 125 ml of NS with 5% BSA as a Control at Time 0 h (referred to as ‘0 h’ in the results). Then, either the middle or lower lobe was cannulated and 10^8^ CFU of PA103 strain in 25 ml of NS with 5% BSA was instilled. No blood was added to the perfusate. After 4 h, a BAL was performed in the injured middle or lower lobe using 125 ml of NS with 5% BSA (referred to as ‘4 h’ in the results). The BAL fluid was centrifuged at 500g for 5 min to pellet cells, and the supernatant was collected and stored at −80°C for subsequent analyses.

### Assessment of functional complement activity in mouse BAL

We adapted a previously published serum assay, with modifications for BAL analysis as reported by us (Sahu et al., 2023). Enzyme-linked immunosorbent assay (ELISA) plates (96-well flat-bottom; #3855, Thermo Fisher Scientific) were coated with LPS (2 μg per well in 100 μl; #L2762, Sigma-Aldrich) and incubated overnight at 4°C. Plates were then washed twice with 0.05% Tween 20 in PBS, followed by a single wash with Mg²⁺-EGTA buffer to prepare for sample addition.

Samples (serum at 1:10 dilution and BAL at 1:5 dilution in Mg²⁺-EGTA buffer) were added at 50 μl per well and incubated at 37°C for 1 h. Plates were washed again, three times, using the same buffer sequence as before. Goat anti-mouse C3 antibody (100 μl per well; #55463, MP Biomedicals), diluted 1:4000 in 1% bovine serum albumin (BSA; #A7906, Sigma-Aldrich) in PBS, was then added and incubated for 60 min. After another three washes, horseradish peroxidase–conjugated donkey anti-goat IgG antibody (100 μl per well, diluted 1:2000 in 1% BSA/PBS; #705-035-147, Jackson ImmunoResearch Laboratories) was applied and incubated for 1 hour. The plates were then washed three times, and TMB Color Substrate (#DY999, R&D Systems) was added (100 μl per well) and incubated at room temperature (RT) for 10 min. To stop the reaction, 1 M sulfuric acid (50 μl per well; #DY994, R&D Systems) was added, and optical density was measured at 450 nm using an Epoch Microplate Spectrophotometer (BioTek).

### Cytokine and chemokine measurements

Serum was collected from mice before and after infection as described above, while BAL was obtained from separate groups of uninfected and infected mice at various time points, as the BAL procedure is terminal. BAL protein levels were quantified using a bicinchoninic acid (BCA) assay. Fresh or thawed BAL samples were diluted (1:10 to 1:50 in PBS) to ensure readings fell within the recommended standard curve range specified by the Pierce BCA Protein Assay Kit (#23227, Thermo Fisher Scientific). Total protein levels were then measured according to the kit’s protocol.

To assess cytokine levels in mouse BAL fluid, samples were diluted 1:2 and analyzed with a multianalyte assay (MILLIPLEX MAP Mouse Cytokine/Chemokine Magnetic Bead Panel, MCYTOMAG-70K-10, Millipore Sigma), following the manufacturer’s instructions and read using a Bio-Rad Luminex 100 multiplex system.

### Full-length C3 ELISA

To quantify total C3 levels in mouse biological samples, we developed a full-length C3 sandwich ELISA. For use with BAL and serum, 96-well flat-bottom ELISA plates (#3855, Thermo Fisher Scientific) were coated with a C3b capture antibody (1:50 dilution; 100 μl per well in PBS; #HM1045, rat anti-mouse C3b/iC3b/C3d/C3dg, clone 11H9; Hycult Biotech) and incubated overnight at 4°C. The plates were then washed three times with PBS containing 0.05% Tween 20 and blocked with 1% bovine serum albumin (BSA; #A7906, Sigma-Aldrich) for 1 h at RT. Following blocking, plates were washed again, and 100 μl of samples (diluted 1:10 for serum and 1:5 for BAL in 1% BSA/PBS) or a standard curve (purified mouse C3; #M113, Complement Technology Inc.) ranging from 400 to 6.25 ng/ml were added to each well. The samples and standards were incubated for 2 h at RT. Plates were then washed three times before adding 100 μl per well of Mouse C3a Detection Antibody (1:1000 dilution in 1% BSA/PBS; Biotin Rat Anti-Mouse C3a, #558251, BD Pharmingen) for 1 h at RT.

After another set of three washes, samples were incubated with 100 μl per well of HRP-conjugated streptavidin (#DY998, R&D Systems) at a 1:200 dilution for 30 min at RT. Following three additional washes, 100 μl of TMB Color Substrate (#DY999, R&D Systems) was added to each well and incubated for 10 min at RT. The reaction was stopped by adding 50 μl per well of 1 M sulfuric acid (#DY994, R&D Systems), and absorbance was measured at 450 nm using an Epoch Microplate Spectrophotometer (BioTek).

### C3a ELISA

The protocol for detecting C3a-neo in supernatant samples was adapted from a previously published serum protocol (Pagano et al., 2009). ELISA plates (96-well flat-bottom; #3855, Thermo Fisher Scientific) were coated with a Mouse C3a Capture Antibody (1:250 dilution; 100 μl per well in PBS; Purified Rat Anti-Mouse C3a, Cat# 558250, BD Pharmingen) and incubated overnight at 4°C. Following this, plates were washed three times with PBS containing 0.05% Tween 20 and then blocked with 1% bovine serum albumin (BSA; #A7906, Sigma-Aldrich) at RT for 1 h. After blocking, plates were washed again, and diluted samples (BAL 1:25, serum 1:50 in 1% BSA/PBS) were added to each well (100 μl per well). A standard curve prepared from purified mouse C3a (Purified Mouse C3a Protein, Cat# 558618, BD Pharmingen) was used, ranging from 50 to 3.125 ng/ml. Samples and standards were incubated for 2 h at RT. The plates were washed three more times, and a Mouse C3a Detection Antibody (1:1000 dilution; 100 μl per well in 1% BSA/PBS; Biotin Rat Anti-Mouse C3a, #558251, BD Pharmingen) was added for 1 h at RT. After another series of washes, 100 μl of HRP-conjugated streptavidin (#DY998, R&D Systems) was added at a 1:200 dilution and incubated for 30 min at RT. Following three additional washes, 100 μl of TMB Color Substrate (#DY999, R&D Systems) was added per well and incubated for 10 min at RT. The reaction was stopped with 50 μl of 1 M sulfuric acid (#DY994, R&D Systems), and absorbance was measured at 450 nm using an Epoch Microplate Spectrophotometer (BioTek).

### Measurement of complement analytes in human BAL

Complement analytes such as C3 and iC3b were quantified in the BAL obtained from the ex vivo human lungs using the MILLIPLEX Human Complement Panel 2 - Immunology Multiplex Assay (HMCP2MAG-19K). Samples were diluted at 1:10 using the assay buffer, and the assay was performed according to the manufacturer’s instructions. Data was acquired using the Luminex 200 system (Thermo Fisher, xPonent software, version 3.1) and analyzed using the Belysa Immunoassay Curve Fitting Software (Version 1.2.1, Sigma-Aldrich).

### IgM ELISA

Levels of IgM in BAL samples from infected mice were measured using the Mouse IgM Uncoated ELISA Kit (Cat# 88-50470-22, Invitrogen). Samples were diluted 1:4000 in 1% BSA/PBS, and the assay was performed according to the manufacturer’s instructions. Levels of IgM in the ex vivo human lung BAL specimens were measured using Human IgM Uncoated ELISA Kit (Cat#88-50620-22, Invitrogen). Samples were diluted 1:4 in assay buffer, and the assay was performed according to the manufacturer’s instructions.

### Total protein quantification in BAL

To quantify BAL protein levels using the bicinchoninic acid (BCA) assay, fresh or thawed BAL samples were diluted 1:50 in PBS to align with the assay’s recommended standard curve range. Total protein concentration was then measured using the Pierce BCA Protein Assay Kit (#23227, Thermo Fisher Scientific), following the manufacturer’s protocol.

### Immunoblotting for C3

Mouse-derived samples were diluted in PBS at a ratio of 1:10 for serum and 1:5 for BAL. Laemmli sample buffer (#1610747, Bio-Rad, Hercules, CA) was added to each sample, and the mixtures were heated for 5 min at 100°C in a heat block to ensure complete protein denaturation. Immunoblot analysis was conducted using samples separated on reduced or non-reduced 4%–20% SDS-PAGE gels (unless stated otherwise), transferred onto nitrocellulose membranes, and probed with specific primary antibodies: goat anti-C3 (#55444, MP Biomedicals 1:3000). Peroxidase AffiniPure™ Donkey Anti-Goat IgG (H+L) (#705-035-147, Jackson ImmunoResearch Laboratories, 1: 3000) secondary antibody was used for detection, and signals were developed with SuperSignal West Pico PLUS chemiluminescent substrate (#34580, Thermo Fisher Scientific). Images were captured using an iBright™ CL1500 Imaging System (A44114, Thermo Fisher Scientific).

### Flow cytometry

To generate a single-cell suspension, mouse lungs were digested with a solution containing 1.5 mg/ml collagenase A (#10103586001, Roche), 0.1 mg/ml deoxyribonuclease I (DNase I; #D4527, Sigma-Aldrich), 5% fetal bovine serum, 10 mM HEPES buffer (#25-060-CI, Corning), and PBS. The lungs were initially inflated with 1 ml of the digestion solution, then cut into 3- to 4-mm sections, and gently vortexed in an additional 2 ml of the solution. Samples were incubated in a shaker incubator at 37°C and 190 rpm for 40 min, with gentle vortexing every 5 to 7 min to enhance tissue breakdown. After incubation, 10 ml of ice-cold PBS was added to halt the digestion, and the suspension was vortexed for 30 sec. The cell mixture was filtered through a 70-μm strainer (#431751, Corning) and centrifuged at 500g for 10 min at 4°C. The resulting supernatant was discarded, and the cell pellet was resuspended in 2 ml of ACK Lysing Buffer (#A10492-01, Gibco) to lyse red blood cells. This suspension was incubated at RT for 4 to 7 min, then diluted in 10 ml of ice-cold PBS containing 1% BSA, and centrifuged again at 500g for 10 min at 4°C. A final cell counts of 1-2 × 10^6^ was transferred into a 96-well plate for antibody staining with FITC anti-mouse CD64 (FcγRI) Antibody (Biolegend#139316, 1:100), PerCP/Cyanine5.5 anti-mouse Ly-6C Antibody (Biolegend#139316, 1:100), APC anti-mouse/human CD11b Antibody, (Biolegend#101212, 1:100), APC/Cyanine7 anti-mouse Ly-6G Antibody (Biolegend#127624, 1:100), MHC Class II (I-A/I-E) Monoclonal Antibody (M5/114.15.2), Super Bright™ 600 (eBioscience, #63-5321-82, 1:1000), Brilliant Violet 711™ anti-mouse CD11c Antibody (Biolegend#117349, 1:100) and PerCP/Cyanine5.5 anti-mouse CD45 Antibody (Biolegend#103131, 1:200). Data acquisition was performed on an Attune NxT flow cytometer (Thermo Fisher Scientific), and FlowJo v10.10.0 (Becton, Dickinson and Company) was used for data analysis.

### Lung immunostaining

Lung sections were deparaffinized, rehydrated in PBS, and subjected to antigen retrieval by heating in Tris-Based Antigen Unmasking Solution (H3301, Vector Laboratories) using a pressure cooker (Decloaking Chamber, Biocare Medical) for 10 minutes. Non-specific binding was blocked with 5% donkey serum and 3% BSA for 30 min at RT. To identify epithelial cells, sections were incubated with primary antibodies diluted in blocking buffer: Rabbit anti-mouse TTF-1 (#WRAB-1231, Seven Hills Bioreagents, 1:1000) overnight at 4°C followed by Donkey anti-Rabbit IgG (H+L) Highly Cross-Adsorbed Secondary Antibody, Alexa Fluor™ 488 (# A-21206, Thermo Fisher Scientific, 1:300) for 60 min at RT. To identify neutrophils, slide was incubated with Ly-6G Monoclonal Antibody (#16-9668-82, Thermo Fisher Scientific, 1:50) overnight at 4°C followed by Goat anti-Mouse IgG (H+L), Cross-Adsorbed, Cross-Adsorbed Secondary Antibody, Alexa Fluor 647 (# A-21235, Thermo Fisher Scientific, 1:300) for 60 min at RT. Nuclei were counterstained with DAPI using ProLong Gold with DAPI (Thermo Fisher Scientific). Fluorescence images were captured with a Leica DM5000B microscope and DFC7000T camera, controlled by Leica Application Suite X software. Brightness and contrast adjustments were performed using ImageJ.

### Drug administration

CXCL2 agonist: Recombinant Mouse CXCL2/MIP-2 Protein (45-M2-010, R&D) was administered orotracheally at a dose of 0.2 microgram/mouse prior to the *Pa* infection, as has previously been reported in models of pneumonia (Fulton et al., 2002). The same volume of PBS, the diluent vehicle was administered as a control.

CXCR2 antagonist: A selective, non-peptide CXCR2 antagonist, SB225002 (AAJ65387MA, Thermo Fisher Scientific) was administered intraperitoneally at a dose of 15 microgram/mouse as two separate doses – one the day before infection and a second dose given again just prior to the *Pa* infection, based on a review of prior literature (Herbold et al., 2010; Cao et al., 2018). The same volume of vehicle control (10% DMSO in corn oil) was administered as a control.

C5aR1 antagonist: PMX53 (CAS 219639-75-5, Sigma-Aldrich) was administered orotracheally at a dose of 3 mg/kg on the same day prior to orotracheal *Pa* infection. The drug was resuspended in DMSO and further diluted in water. Control mice received a dilution of DMSO in water.

### RNA extraction and sequencing

Total RNA was extracted from lung lobes using the RNeasy Plus Mini Kit (Cat# 74134, QIAGEN). Tissues were placed in RNA*later*^TM^ stabilization solution (Cat# AM7020, Thermo Fisher Scientific) and stored as instructed by reagent’s protocol. Stored lungs were homogenized with the kit reagents and processed according to the manufacturer’s instructions. Extracted RNA was then submitted to the Genome Technology Access Center at the McDonnell Genome Institute (GTAC@MGI), Washington University School of Medicine for RNA sequencing. Ribosomal depletion, RiboErase (HMR) was employed for library preparation and sample processing. Sequencing was performed on a NovaSeq S4 platform with a target of at least 30 million reads per library using 2×150 paired-end reads. Dual indexing was used during library construction to ensure unique sample identification.

### Bulk-RNA sequencing analysis

Samples were prepared according to library kit manufacturer’s protocol, indexed, pooled, and sequenced on an Illumina NovoSeq 6000. Basecalls and demultiplexing were performed with Illumina’s bcl2fastq software and a custom python demultiplexing program with a maximum of one mismatch in the indexing read. Fastq files was uploaded to Partek Flow Software (Illumina) for analysis. Adapter trimming and quality filtering were performed. RNA-seq reads were then aligned to the mouse reference genome (mm10) using the STAR aligner version 2.7.8a (Dobin et al., 2013). Gene-level quantification was performed in Partek Flow to generate read count matrices. Sequencing performance was assessed for the total number of aligned reads, total number of uniquely aligned reads, and features detected. Low-expression features were filtered out, excluding those with a maximum count of ≤50. Principal component analysis (PCA) was performed to identify and remove outliers, resulting in a final sample size of n=4 per group. Samples were grouped by genotype (*C3^f/f^Scgb1a1CreER^T2^*^+/-^ vs. *C3f/f (i.e. C3^f/f^Scgb1a1CreER^T2^*^-/-^)) and differential gene expression analysis was conducted between the groups. Genes were filtered based on a false discovery rate (FDR) step-up threshold of ≤0.05. A heatmap and volcano plot were generated using the filtered gene list to visualize differential expression patterns. Pathway analysis was performed using EnrichR to identify common biological themes among the significantly downregulated genes from the differential gene expression analysis. The gene list was uploaded to EnrichR, and the most significant ontologies were identified using the GO Biological Process 2023 database. To complement this approach, an independent gene-ontology analysis was conducted using COMPBIO (Comprehensive Multi-omics Platform for Biological InterpretatiOn), a biological knowledge generation platform as done previously (Sahu et al., 2023).

Downstream analysis was conducted in R (version 4.4.2) using the *DESeq2* package (version 1.46.0). The data consisted of paired samples. Lowly expressed genes were filtered out, and count data were normalized using DESeq2’s median-of-ratios method. Principal component analysis (PCA) was used to assess sample variation. Differential expression analysis was performed using DESeq2, with p-values adjusted using the Benjamini–Hochberg method. Genes with an adjusted p-value < 0.05 and absolute log2 fold change ≥ 1 were considered significantly differentially expressed. For gene set enrichment and pathway analysis, we employed the EnrichR web-based tool and the GSEA algorithm, in conjunction with several R packages. We used *org.Mm.eg.db* to convert Ensembl gene IDs to gene symbols and performed enrichment analyses using the *clusterProfiler* and *ReactomePA* packages. Specifically, we applied the enrichGO and enrichKEGG functions to identify enriched Gene Ontology (GO) terms and KEGG pathways, respectively. Visualization of results, including PCA plots, heatmaps, and volcano plots, was performed using the *ggplot2* and *pheatmap* packages in R.

### Single-cell RNA-sequencing

cDNA was prepared after the GEM generation and barcoding, followed by the GEM-RT reaction and bead cleanup steps. Purified cDNA was amplified for 11-13 cycles before being cleaned up using SPRIselect beads. Samples were then run on a Bioanalyzer to determine the cDNA concentration. GEX libraries were prepared as recommended by the Chromium Next GEM Single Cell 3’ HT Reagent Kits v3.1 (Dual Index) (CG000416) with appropriate modifications to the PCR cycles based on the calculated cDNA concentration. For sample preparation on the 10x Genomics platform, the Chromium Next GEM Single Cell 3’ HT Kit v3.1 - 48 rxns (PN-1000348), Chromium Next GEM Chip M Single Cell Kit-| 16 rxns (PN-1000371) and Dual Index Kit TT Set A, 96 rxns (PN-1000215) were used. The concentration of each library was accurately determined through qPCR utilizing the KAPA library Quantification Kit according to the manufacturer’s protocol (KAPA Biosystems/Roche) to produce cluster counts appropriate for the Illumina NovaSeq X Plus instrument. Normalized libraries were sequenced on a NovaSeq X Plus Flow Cell using the151×10×10×151 sequencing recipe according to manufacturer protocol. Read 1 was trimmed to the 10x Genomics recommendation of 28bp. A median sequencing depth of 50,000 reads/cell was targeted for each Gene Expression Library.

Sequencing data were processed using the CellRanger pipeline (version 8.0.1) with default settings, aligning reads to the Mouse mm10 (GENCODE vM23/Ensembl98) reference genome. The resulting gene expression matrix was first processed with SoupX to remove ambient RNA contamination. The cleaned matrix was then used for downstream bioinformatics analysis. The Seurat package (version 5.1.0) in R (version 4.4.2) was used for data processing and analysis. Cells with at least 500 detected genes and mitochondrial content below 25% were retained for further steps. Normalization and variance stabilization were performed using the SCTransform method in Seurat, and a few genes were excluded from the variable feature selection step. Thirty principal components were used to carry out dimensionality reduction, followed by Louvain clustering and Uniform Manifold Approximation and Projection (UMAP) visualization. Cell type-specific markers were identified with Seurat’s FindAllMarkers function, with log fold change set to 0.25. Annotation for broad cell types was performed using known markers, and subtypes were further identified with specific gene markers. Differentially expressed genes between clusters were determined using the Wilcoxon rank sum test with default parameters in Seurat.

For the analysis of cell–cell communication, we utilized CellChat (version 2.1.2), which consists of two main components: the Ligand-Receptor Interaction Explorer, which allows for efficient exploration of ligand-receptor interactions, and the Cell-Cell Communication Atlas Explorer, which facilitates the analysis of cell-cell communication patterns in scRNA-seq datasets processed (Jin et al., 2025). This user-friendly R toolkit, along with its web-based platform (http://www.cellchat.org/), enables the identification of intercellular signaling pathways and the construction of cell–cell communication networks. In our analysis, we specifically selected PMNs (neutrophils) as the target cell type, and type 2 alveolar epithelial cells (AT2), alveolar macrophages (AM), and Myofibroblasts/SMCs (MyoFB/SMC) as source cell types, focusing on the inferred interactions between these cell populations of interest.

### Statistical analyses

Data are presented as scatterplots showing individual data points.Dispersion is shown by mean ± SD. Unpaired two-tailed t test was used for comparing two groups. Ordinary one-way analysis of variance (ANOVA) with multiple comparisons testing was used for more than two groups. A P value of <0.05 was considered significant, although values of >0.05 have been reported when there was evidence of a trend. Prism v10.3.0 (GraphPad) was used for statistical analysis.

### Data availability statement

The sequencing data has been deposited on NCBI GEO and will be made publicly available at the time of publication.

## ACKNOWLEDGEMENTS

The authors thank Angel Lu and Aayusha Thapa for their technical assistance, and Drs. Maxwell Sanfilippo-Burchman, Alexander Earhart, John Michael Sanchez, Jungheun Hyun, Xiaobo Wu and John P. Atkinson for their inputs. We thank the Genome Technology Access Center at the McDonnell Genome Institute at Washington University School of Medicine for help with genomic analysis and accurate reporting of the sequencing methods. The Center is partially supported by NCI Cancer Center Support Grant #P30CA91842 to the Siteman Cancer Center from the National Center for Research Resources (NCRR), a component of the National Institutes of Health (NIH), and NIH Roadmap for Medical Research. This publication is solely the responsibility of the authors and does not necessarily represent the official view of NCRR or NIH. We thank members of the Immune Assessment Core (Monica Cappelletti, Itzel Ramirez and Cache Robinson) for their assistance with multiplex ELISAs.

