## Supplementary Figure File for "Kinetics of local C3 production orchestrates neutrophil recruitment in lung injury"

### Supplementary Figure S1

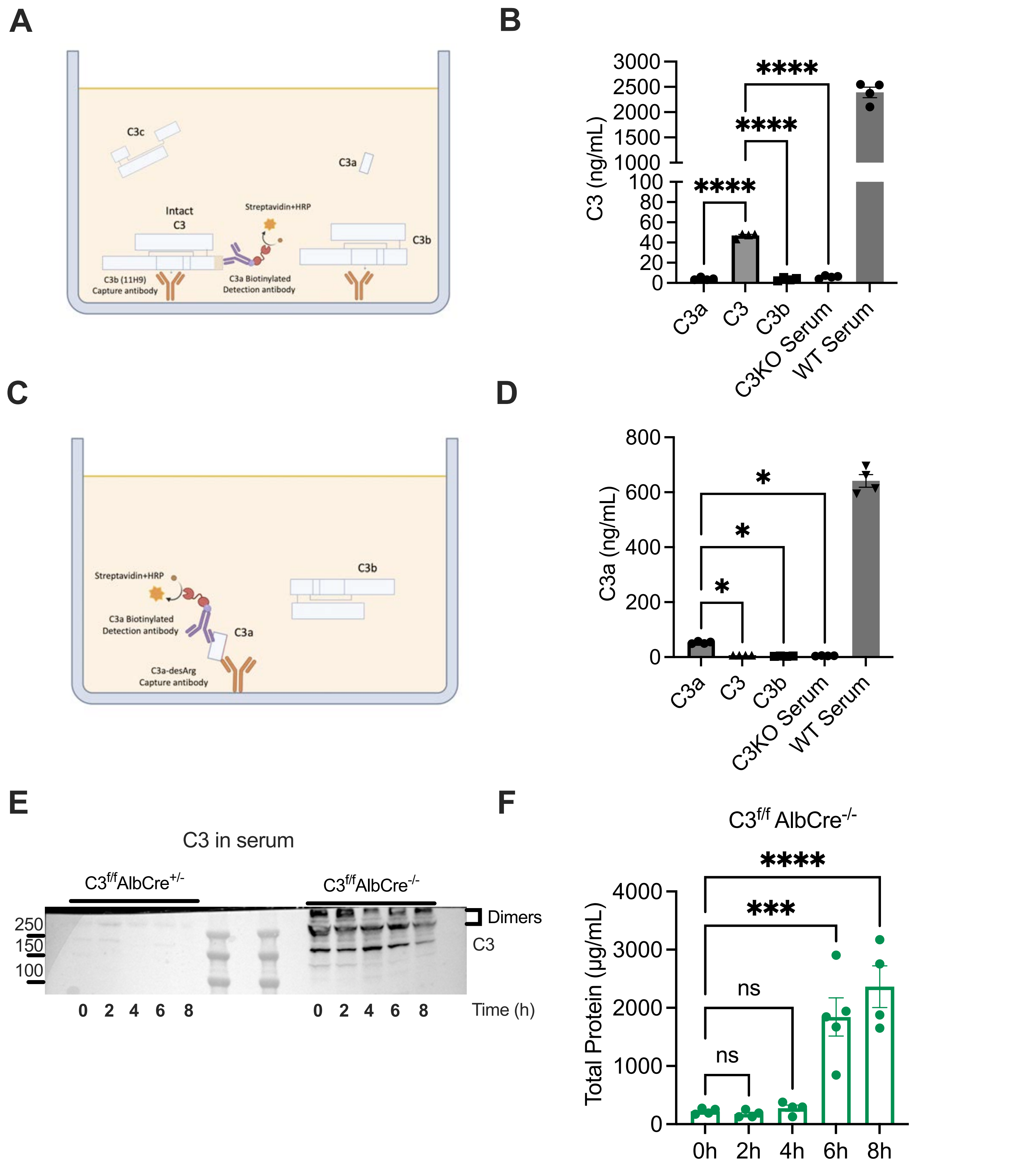


**Figure S1. Circulating C3 requires an ongoing infection to leak into the bronchoalveolar space.** Purified proteins and serum from wildtype (WT) mice used to confirm ELISAs detecting (**A & B**) full-length/intact C3 and (**C & D**) cleaved C3a neo-epitope. **(E)** Validation of $C3^{f/f}$ $AlbCre^{+/-}$ mice via non-reducing western blot for serum drawn at multiple time points post-*Pa* infection. (**E**) Protein levels of $C3^{f/f}$ $AlbCre^{-/-}$ mice by BCA assay. Distribution of data is represented as scatterplots showing individual data points, and bars represent means ± SD. ****p<0.0001, ***P < 0.001, **P < 0.01, and *P < 0.05 using unpaired two-tailed t test.

### Supplementary Figure S2

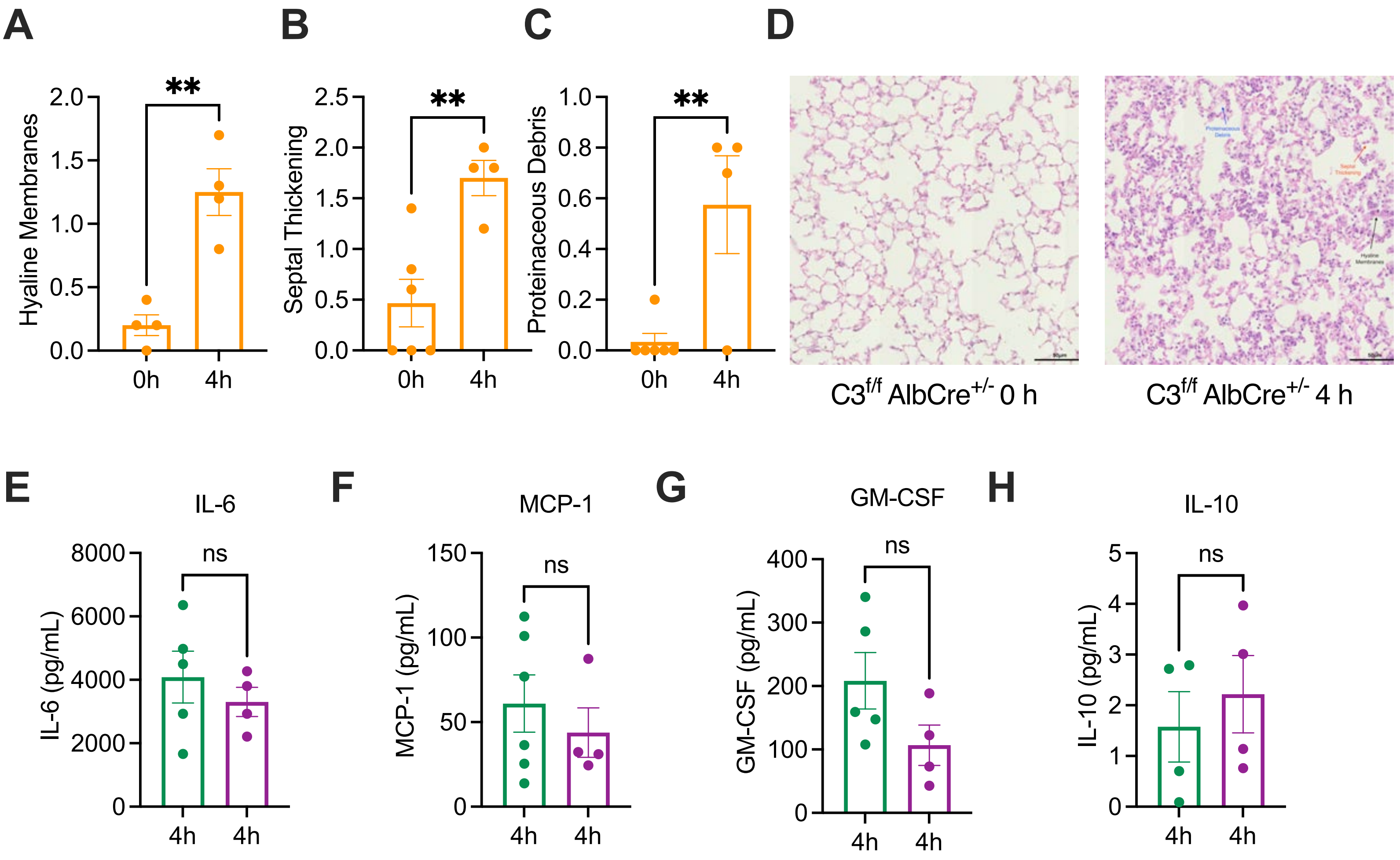


**Figure S2. Lung injury parameters and cytokine levels at 4 h post-infection. (A-D)** Parameters for lung injury at 4h after *Pa* infection in C3 liver-deficient mice (orange), including lung scoring for **(A)** hyaline membrane, (**B**) septal thickening and (**C**) proteinaceous debris. (**D**) Representation of parameters in H&E staining for $C3^{f/f}$ $AlbCre^{+/-}$ Mice (Black: Hyaline Membrane, Red: Septal Thickening and Blue: Proteinaceous Debris). Multiplex ELISA for (**E**) IL-6, (**F**) MCP-1, (**G**) GM-CSF and (**H**) IL-10 in the bronchaolveolar lavage fluid of $C3^{f/f}Scgb1a1CreER^{T2-/-}$ (green) and $C3^{f/f}Scgb1a1CreER^{T2+/-}$ (purple) mice at 4 h post infection. Distribution of data is represented as scatterplots showing individual data points, and bars represent means ± SD. $^{**}P < 0.01$, and $^{*}P < 0.05$ using unpaired two-tailed t test.

### Supplementary Figure S3

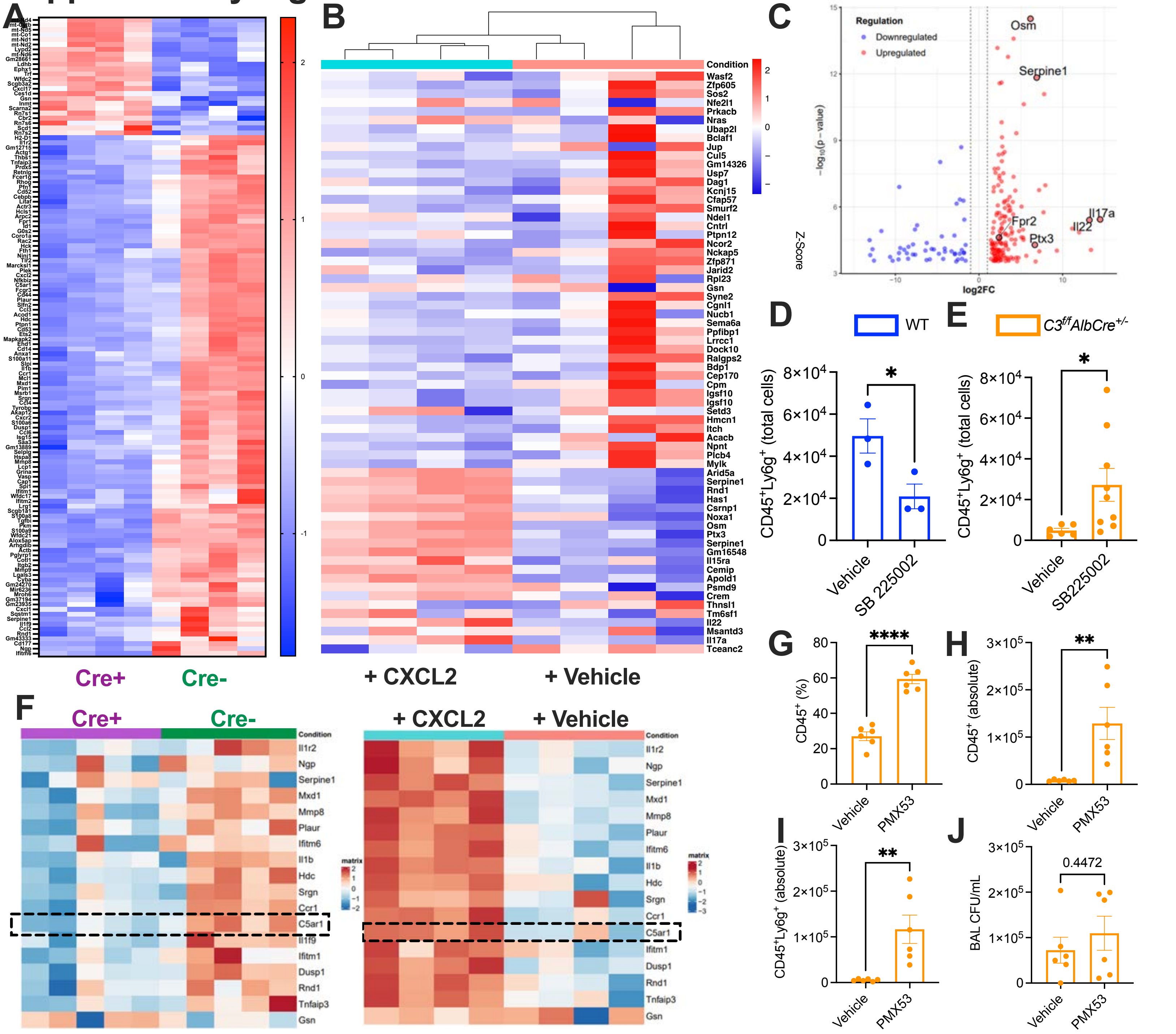


**Figure S3. C3-mediated neutrophil recruitment to the lung in the initial hours of a bronchopulmonary infection occurs through multiple pathways.** (**A**) Heat map comparing 130 genes obtained from differential analysis expression and filtered by FDR step up ≤0.05 from lungs of mice deficient in lung epithelial cell-derived C3 ($Cre^+$) compared to controls ($Cre^-$) at 4 h post-*Pa* infection. (**B**) Similar heatmap representing differential gene analysis from the lungs of mice deficient in lung epithelial cell-derived C3 given orotracheal CXCL2 vs. vehicle control, at 4 h post-*Pa* infection. Restricted to 72 genes above logFC threshold of -5 to +5. (**C**) Volcano plot of all 229 genes from comparison in B, with log2FC on x-axis and p value (-log10) on y-axis. (**D**) Total BAL neutrophils ($CD45^+$, $Ly6g^+$) measured using flow cytometry at 4 h post-*Pa* infection in lungs of wildtype mice injected intraperitoneally (IP) with SB225002. (**E**) Total BAL neutrophils ($CD45^+$, $Ly6g^+$) measured using flow cytometry at 4 h post-*Pa* infection in whole lung of liver-deficient C3 mice (yellow) injected intraperitoneally (IP) with SB225002. (**F**) Differently expressed genes common to both datasets in S3A and S3B/C. (**G-J**) Liver-deficient C3 mice treated oropharyngeally with PMX53, infected with *Pa* and euthanized at 4 h post-infection. Percentage (**G**) and absolute number of $CD45^+$ cells (**H**) and neutrophils (**I**) in BAL. (**J**) Colony forming units (CFU) of bacteria in BAL. Distribution of data is represented as scatterplots showing individual data points, and bars represent means ± SD. *P < 0.05, **P<0.01, ****P<0.00001 using unpaired t-test.
